# Deletion of *Gremlin-2* alters estrous cyclicity and disrupts female fertility in mice

**DOI:** 10.1101/2020.10.01.322818

**Authors:** Robert T. Rydze, Bethany Patton, Hannia Salazar-Torralba, Shawn Briley, Gregory Gipson, Rebecca James, Aleksandar Rajkovic, Thomas Thompson, Stephanie A. Pangas

## Abstract

Members of the differential screening-selected gene aberrative in neuroblastoma (DAN) protein family are developmentally conserved extracellular binding proteins that antagonize bone morphogenetic protein (BMP) signaling. This protein family includes the Gremlin proteins, GREM1 and GREM2, which are known to have key functions during embryogenesis and adult physiology. While BMPs play essential roles in adult female reproductive physiology, the role of the DAN family in ovarian function is less understood. We generated mice null for *Grem2* to study its role in female fertility in addition to screening patients with primary ovarian insufficiency for variants in GREM2. *Grem2^-/-^* mice are viable and female *Grem2^-/-^* mice have diminished fecundity and irregular estrous cycles. This is accompanied by reduced serum levels of anti-Müllerian hormone, a marker of the ovarian reserve, in adult mice. Alterations in ovarian expression of inhibin and activin subunit genes, which are required for regulation of the hypothalamic-pituitary-ovarian (HPO) axis, were identified. While *Grem2* mRNA transcript was not detected in the pituitary, *Grem2* was expressed in the hypothalami of wild type female mice. Additionally, screening 106 women with primary ovarian insufficiency identified one individual with a heterozygous variant in GREM2 that lies within the predicted BMP-GREM2 interface. In total, these data suggest that *Grem2* is necessary for female fecundity by playing a novel role in regulating the HPO axis and possibly contributing to female reproductive disease.

## Introduction

Gremlin-2 (GREM2) is a member of the *differential screening-selected gene aberration in neuroblastoma* (DAN) family of bone morphogenetic protein (BMP) antagonists and is also known as *protein related to DAN and Cerberus* (PRDC) (1–3). It was originally identified in a gene trap for developmentally important genes (4). The DAN family includes GREM2/PRDC, Gremlin-1 (GREM1), Sclerostin, DAN, Cerberus, Caronte, Coco, and Dante (5). This protein family is best characterized as extracellular binding proteins that sequester BMPs, thereby preventing them from binding and activating their signaling receptor complex. GREM2 strongly inhibits BMP signaling with high affinity for BMP2, BMP4, and BMP7 (6). GREM2 also binds other TGFβ family members, including anti-Müllerian hormone (AMH) and with low affinity to GDF5 (6,7). GREM2 tightly associates with heparin through a protein binding domain outside of the BMP binding domain, which limits or downregulates BMP signaling, thus enhancing the inhibitory activity of GREM2 on BMPs (2,8).

Human genome-wide association studies (GWAS) identified *GREM2* variants associated with developmental disorders and disease, such as bone function and bone mass associated with osteoporosis, atrial fibrillation, and tooth agenesis (9–13). A *Grem2^-/-^* mouse line was previously developed as part of a high-throughput mouse knockout and phenotyping project (14). Skeletal phenotypes such as elevated spine and femur bone mineral density as well as tooth defects were identified as the major defects in *Grem2^-/-^* mice (14). No fertility defects were reported in male or female *Grem2^-/-^* mice, although the reproductive phenotyping screen was limited (P. Vogel, personnel communication). A variant of *GREM1* has been reported in a patient with primary ovarian insufficiency (POI), but none have been reported for *GREM2* (15).

There are limited studies characterizing GREM2 function in female reproductive biology. *GREM2* is expressed in the developing human ovary at 8-21 weeks gestation, with increasing expression towards the time of primordial follicle formation (16). Embryonic *Grem2* expression in the developing mouse gonad (male or female) has not been not characterized, but *Grem2* is expressed in the mouse and rat ovary in granulosa cells during postnatal follicle development (1). *In vitro* studies indicate GREM2 inhibits BMP4 and AMH, both of which regulate growth dynamics of the primordial to primary follicle transition, although in opposing directions (*i.e.*, BMP4 promotes while AMH inhibits the transition) (1,7,17–19), possibly indicating a role for GREM2 in regulating growth dynamics of the ovarian reserve.

Given above studies implicating GREM2 in ovarian folliculogenesis, we generated a new knockout mouse model of *Grem2* to determine its role in mammalian reproduction. We found that *Grem2^-/-^* females have altered fecundity due to changes in reproductive cyclicity and a reduction of the ovarian reserve marker, AMH, while not grossly affecting folliculogenesis. We additionally identified a heterozygous novel nonsynonymous mutation in a patient with POI. These data suggest that GREM2 may be required to regulate reproductive function in both mouse and human, possibly at multiple levels within the HPO axis.

## Materials and Methods

### Generation of *Grem2^-/-^* mice

Experimental animals were used in accordance with the National Institutes of Health Guide for the Care and Use of Laboratory Animals using Institutional Animal Care and Use Committee approved protocols at Baylor College of Medicine. Mice were maintained on a *C57BL/6/129S7;SvEvBrd* mixed hybrid background, which was the genetic background to our previous knockout of *Grem1* (20) as well as our other published lines. Mice were housed in microisolator cages with acidified water on a fixed 12-hr light and 12-hour dark cycle and fed ad libitum on a breeder rodent chow (5053 PicoLab Rodent Diet 20, Richmond, IN) supplemented twice weekly with a soft pellet dietary supplement (LoveMash Rodent Reproductive Diet, Bio-Serve, Flemington, NJ). Incisor length and weight was monitored during their housing.

A *Grem2* null allele was generated at the Embryonic Stem Cell and Genetically Engineered Mouse Cores at Baylor College of Medicine. Single guide RNA (sgRNA) sequences were selected to flank the genomic region surrounding exon 2 of the *Grem2* gene (upstream sgRNA 5’-GGGGTAGATGGTGCTACTTC **CGG;** downstream sgRNA, 5’-GAAAAATCTTGTCGAGTTTC **TGG;** PAM sequences are in bold) using the CRISPR Design Tool (21) and examined for potential off target mutagenesis. Neither guide was predicted to have off target effects. DNA templates for *in vitro* transcription of sgRNAs were produced using overlapping oligonucleotides (in a high-fidelity PCR reaction (22) and sgRNA was transcribed using the MEGA shortscript T7 kit (ThermoFisher, Waltham, MA). Cas9 mRNA was purchased from ThermoFisher. The BCM Genetically Engineered Mouse Core microinjected Cas9 mRNA (100 ng/μl) and sgRNA (10 ng/μl) into the cytoplasm of 100 pronuclear stage C57Bl/6J embryos. Cytoplasmic injections were performed using a microinjection needle (1 mm outer and 0.75 mm inner) with a tip diameter of 0.25-0.5 μm, an Eppendorf Femto Jet 4i to set pressure and time to control injection volume (0.5-1 pl per embryo). Injections were performed under a 200-400× magnification with Hoffman modulation contrast for visualizations.

A single PCR reaction using three primers (P1: 5’-TGTTGTTGTTGTTGACAAAATACTTG; P2: 5’-AATACGAGAAAGCCGTGCTG; P3: 5’-AAAGAGGTGGTGGTGTCCAG) identified the wild type allele (251 bp product) and the deletion allele (~ 510-520 bp product) in putative founders and resulting offspring. Deletion of *Grem2* in founder mice was verified by Sanger sequencing of the PCR-amplified deletion allele junction fragment. Two founder mice were initially characterized but no difference in phenotypes (presence of an incisor defect; similar numbers of pups per litter; irregular litter production) were detectable, so one founder line was chosen for more detailed study. Mice were genotyped by PCR of genomic DNA isolated from ear punches or tail snips. To minimize potential off-target mutation effects, founder mice were backcrossed to F1 *C57BL/6/129S7;SvEvBrd* mice prior to establishing homozygous mating to generate *Grem2^-/-^*. Wild type mice of the same mixed background (*C57BL/6/129S7;SvEvBrd*) were used as controls (“wild type”).

### Fertility Studies

Individually housed female mice of each genotype were bred at 6-8 weeks of age continuously to wild type *C57BL/6/129S7;SvEvBrd* F1 hybrid males of known fertility for eight months. The number of pups born at 4-week intervals (one ‘month’) was recorded beginning when each of the mating pairs was set up as individual pair-breeders. For estrous cycle analysis, six-month old mice were individually housed with enrichment (EnviroPak, Lab Supplies, Dallas, TX) and vaginal lavage and cytology was performed daily at 9:00am for 1 month as previously described to identify four stages: estrus, diestrus, proestrus and metestrus (23).

### Histologic Analysis

Mice were weighed prior to necropsy then anesthetized by isoflurane inhalation (Abbott Laboratories, Abbott Park, IL) and euthanized by cervical dislocation. Estrous cycle was determined at time of necropsy, if not already known. Ovaries were collected and fixed in 10% neutral buffered formalin (Electron Microscopy Sciences, Hatfield, PA) overnight, transferred to 70% ethanol, and processed and embedded at the Human Tissue Acquisition and Pathology Core at Baylor College of Medicine using standard protocols. Follicle counts were performed as previously described (24). Briefly, ovaries from 3-week mice were serially sectioned at 5-μm and all sections retained. Sections were stained in periodic acid-Schiff (PAS) (Sigma, St. Louis, MO). Only follicles containing an oocyte with a visible nucleus (primordial, primary, secondary, antral, atretic) were counted in every fifth section to avoid double counting oocytes. Final values of preantral follicles were multiplied by a correction factor of 5 based on previous published methodologies (25). Statistical differences in the total number of follicles were assessed using Student’s *t*-test.

### Hormone Analysis

For hormone analysis, blood was retrieved from deeply anesthetized (isoflurane) diestrous stage mice by cardiac puncture, and serum separated by centrifugation in microtainer collection tubes (SST BD Microtainer, Becton, Dickinson and Company, Franklin Lakes, New Jersey) and stored at −20°C until assayed. Estradiol (ELISA, CalBiotech), follicle stimulating hormone (FSH)/luteinizing hormone (LH) (ELISA, EMD Millipore), AMH (ELISA, Ansh Laboratory), and testosterone (ELISA, IBL) were quantified by the University of Virginia Ligand Core Facility (Specialized Cooperative Centers Program in Reproductive Research NICHD/NIH U54-HD28934). Assay method information is available online (https://med.virginia.edu/research-in-reproduction/ligand-assay-analysis-core/assay-methods/). For statistical analysis, values that fell below the threshold of detection were set to the value for the lower limit of detection (26). As ELISA data are not normally distributed, data were log transformed prior to statistical analysis.

### Immunohistochemistry

Immunohistochemistry was performed as previously described (27) using the Vectastain ABC method (Vector Laboratories, Burlingame, CA) for analysis of the macrophage marker, F4/80 (rat anti-F4/80, catalog #AB6640, Abcam, Cambridge UK, 1:100). Immunoreactivity was visualized by diaminobenzidine (DAB) (Vector Laboratories) and ovaries counterstained in hematoxylin. Fluorescent immunohistochemistry was used to visualize AMH (goat anti-AMH, 1:250; catalog #6886, Santa Cruz Biotechnology, Santa Cruz, CA). Briefly, 5 μm tissue sections were subject to antigen retrieval (0.01 M citric acid and 0.1% Triton X) (Sigma, St. Louis, MO), blocked with the avidin/biotin blocking kit (Vector Laboratories), followed by incubation 3% BSA in Tris-buffered saline (TBS) for non-specific binding. The primary antibody incubated overnight at 4°C in a humidified chamber. Slides were washed in TBS-0.1%Tween (TBS-T) and incubated at room temperature with Alexa Fluor rabbit anti-goat 594 (Invitrogen; 1:250) for 1 hour, washed, incubated in 4’6’-diamidino-2-phenylindole (DAPI) (1:1000) in TBS for 5 minutes then mounted in Vectashield (Vector Laboratories). Fluorescent images were captured using a Nikon A1R-s confocal laser scanning microscope at the BCM Integrated Microscopy Core and processed with the Nikon Perfect Focus System (Nikon Corporation, Japan). Representative follicles within ovary section from wild type and *Grem2^-/-^* mice (n=3 ovaries for each genotype) were analyzed using ImageJ software (ImageJ 1.52a Wayne Rasband, National Institutes of Health, USA http://imagej.nih.gov/ij) to measure the mean fluorescence intensity. Follicle type was based on the classification system developed by Pedersen et al (28). From these values, the statistical difference between wild type and *Grem2^-/-^* follicles was compared using Student’s *t*-test using Graph Pad Prism 5 (GraphPad Software La Jolla, CA).

### Quantitative PCR

Tissues were harvested from diestrous stage females unless otherwise indicated and incubated in RNA*later* (Ambion, Austin, TX) overnight at 4°C then stored at −80°C until use. RNA was isolated using the RNeasy Micro kit (Qiagen, Valencia, CA) with in column DNase treatment (Qiagen) following the manufacturer’s protocol. RNA concentration was quantified using a NanoDrop Spectrophotometer ND-1000 (NanoDrop Technologies, Wilmington, DE). Complementary DNA was synthesized from 200ng of total RNA with the High-Capacity RNA-to-cDNA reverse transcription kit (Life Technologies, Waltham, MA). Real-time quantitative PCR (qPCR) assays were performed on an Applied Biosystems StepOne Realtime PCR System using TaqMan Fast Master mix and predesigned primer-probe mixes for *Grem2* (FAM labeled Mm00501909_m1) and *Gapd* (FAM labeled Mm00484668_m1) or Fast SYBR Green Master Mix (Life Technologies, Waltham, MA) with custom primers chosen from validated qPCR primer sets at <u>PrimerBank</u>. Primer sequences are available upon request. Custom DNA oligonucleotide primer pairs were synthesized at Integrated DNA Technologies (IDT, Coralville, Iowa). Melt curve analysis was used to validate a single amplification peak when using SYBR Green Master Mix. Relative level of transcript was calculated using the ΔΔCT method (29) with the housekeeping gene *Gapd* used for normalization and data shown mean to the relative level in wild type ovaries.

### POI Patient Sample Analysis and Protein Modeling

Whole exome sequence data (WES) were analyzed from two previously published datasets (30,31). These data included 103 patients diagnosed with primary ovarian insufficiency (POI) at the University of Pittsburgh under the approved Institutional Review Board protocols (PRO09080427). Informed consent was obtained from all individual participants. Gene variants were evaluated using the guidelines of the American College of Medical Genetics and Genomics (ACMG), which recognizes five classes of variants: benign, likely benign, uncertain significance, likely pathogenic, and pathogenic (32) and variants with minor allelic frequency in the Exome Aggregation Consortium (ExAC) database > 1% were excluded. Variants not present in the 1000 Genomes Project, Exome Variant Server data sets, Exome Aggregation Consortium (ExAC, Cambridge, MA), or the Single Nucleotide Polymorphism database (dbSNP) were considered novel variants (33–35). Protein modeling was performed as previously described (6) using PyMol (The PyMol Molecular Graphics System, Schrödinger, LLC, New York, NY).

### Statistical Analyses

Statistical analysis was carried out using GraphPad Prism 5 (GraphPad Software, La Jolla, CA). Two-tailed unpaired Student *t*-test using Mann-Whitney U was used for single comparisons. One-way analysis of variance followed by Fisher least significant difference test was used for multiple comparisons. Data not normally distributed (e.g., hormone data and qPCR data) were log transformed prior to statistical analysis. Linear regression was used to assess correlation between FSH and estradiol levels. Sample sizes are indicated in the text and figure legends and a minimum of at least three independent experiments was carried out at all times, with P<0.05 considered statistically significant.

## Results

### Generation and validation of a *Grem2^-/-^* mouse model

A null allele for *Grem2* was engineered using CRISPR/Cas9 genome editing (Fig. 1). Two sgRNAs were designed to target Cas9 to flanking regions of exon 2, generating an approximately one kilobase deletion that contains the splice acceptor site in exon 2, the entire coding region, and portions of the 3’ UTR (Fig. 1A). Exon 1, which encodes the 5’ untranslated region (UTR) and elements of the 3’ UTR remain. sgRNA was injected into pro-nuclear stage embryos along with *Cas9* mRNA. Nonhomologous end joining (NHEJ)-mediated repair of the two double stranded breaks (DSBs) created by sgRNA targeted Cas9 should result in a null allele through loss of exon 2. Seventeen live-born mice were obtained from 100 injected and transferred embryos. PCR genotyping indicated that 4 pups contained a molecular weight band that approximated the predicated size of the deletion (Fig. 1B). Different deletion sizes are produced in each founder because of the imprecise nature of NHEJ and DSB repair. DNA from the four potential founders was sequenced, and two potential founder alleles aligned with the expected deletion (Fig. 1C). These mice were individually crossed to F1 mixed hybrid strain *C57BL/6/129S7/SvEvBrd*, the genetic background for our previous studies on BMP and GREM1 function in the ovary (20,36). Breeding to the wild type strain ensures germline transmission of the correct allele and reduces the probability of carryover of potential off-target mutations. Homozygous mice for both founder lines were generated from heterozygous crosses and were produced at normal Mendelian ratios for both founders. Male and female homozygous mice were viable. No difference in phenotype was detected in initial studies between founder lines (*e.g.*, incisor defects and fertility testing), so the founder line that contained the larger deletion (“P2” Fig. 1B) was chosen for further characterization.

**Fig. 1.**
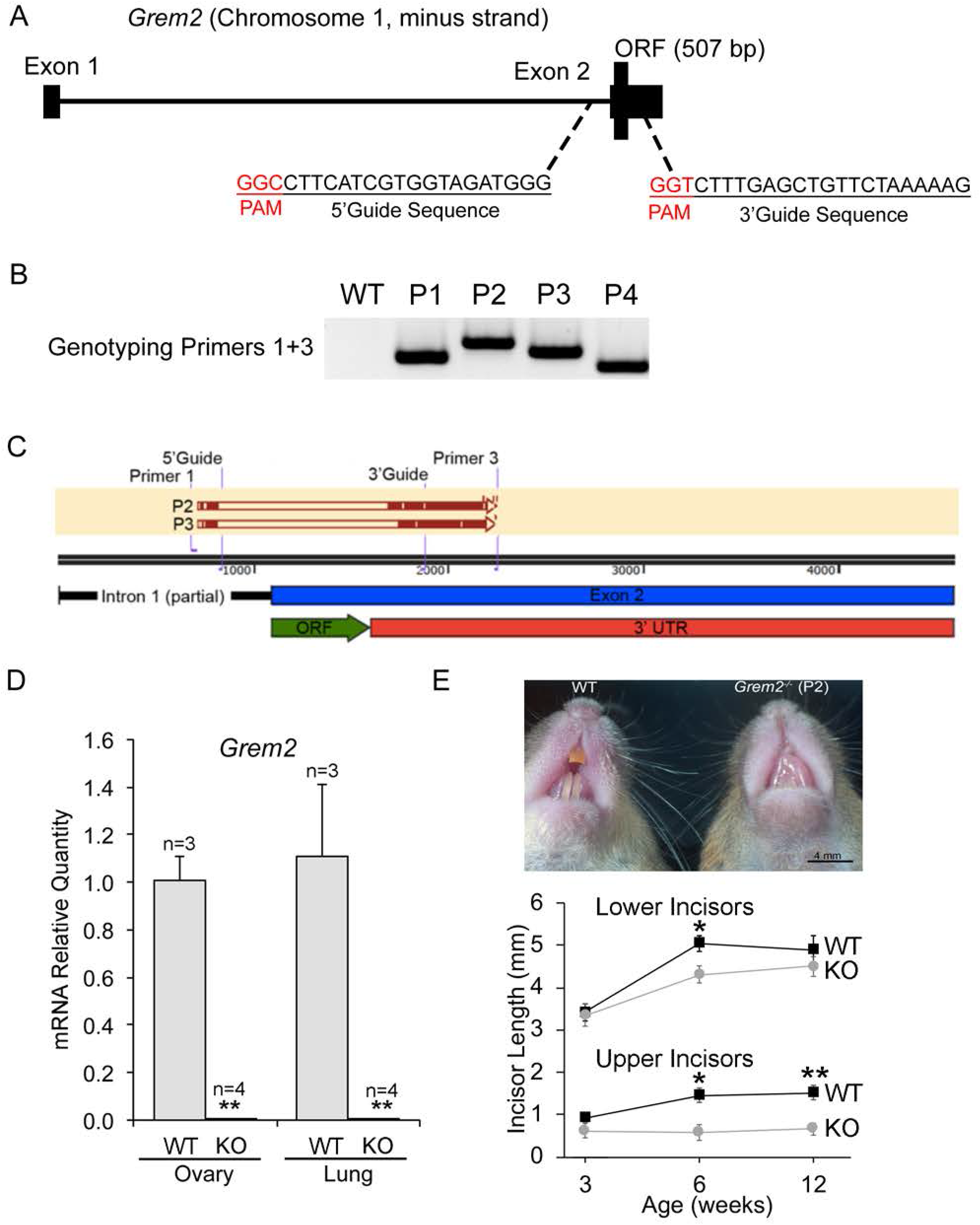
Generation and validation of a *Grem2* null allele. (A) Schematic of the *Grem2* locus on chromosome 1. *Grem2* contains two exons, with the open reading frame (ORF) encoded in exon 2. The 5’ and 3’ guide sequences are shown with the protospacer adjacent motif (PAM) site in red. The 5’ guide sequence is located in intron 1 and the 3’ guide sequence is located within the 3’ untranslated region (UTR). (B) PCR genotyping of genomic DNA of potential founder mice, labeled P1-P4. Different size deletions are typical due to the imprecise nature of NHEJ. (C) Summary of DNA sequencing information and alignment of the founders P2 and P3 with the location of the ORF shown in green and the 3’ UTR in red. (D) Validation of loss of *Grem2* transcript in *Grem2^-/-^* tissues by qPCR, n=3 independent ovaries for wild type (WT) and n=4 for knockout (KO), **P<0.01. Levels are normalized to *Gapd* and shown relative to the amount in the wild type ovary. (E) Comparison of incisors between wild type and *Grem2^-/-^* from the P2 parental line, which was chosen from 2 founder lines with similar fertility defects. Graph shows data for upper and lower incisors in wild type and *Grem2^-/-^* at three ages. No difference was found in upper or lower incisor length between wild type and *Grem2^-/-^* at 3 weeks of age (n=6 mice each genotype). Lower and upper incisors of Grem2^-/-^ (n=5 mice) were significantly smaller at 6 weeks of age (*P<0.05) but only upper incisors were significantly smaller (**P<0.01) in *Grem2^-/-^* (n=5) compared to wild type (n=5) at 12 weeks of age. Image insert, wild type and *Grem2^-/-^* incisors at 9 months of age. Scale bar in photograph, 4mm.

We confirmed loss of *Grem2* expression by measuring *Grem2* mRNA levels by quantitative PCR (qPCR) in tissues known to highly express the transcript, which includes the ovary and lung. As predicted, *Grem2* transcript levels were undetectable in either tissue in *Grem2^-/-^* mice relative to the respective levels in wild type mice (Fig. 1D). As a previous null allele of *Grem2* demonstrated reduced breadth and depth of upper and lower incisors in *Grem2^-/-^* mice over 4 months of age (14), we also measured incisor length in adult animals of our new line. Similar to the previous model, sexually mature (i.e., over 6 weeks of age) *Grem2^-/-^* female mice had defects in incisor length that mainly affected the upper incisors (Fig. 1E). This did not have a major effect on their body weight, as *Grem2^-/-^* females had similar body weights to wild type mice at 3 weeks of age, were slightly but significantly larger than wild type mice at 6 and 12 weeks, but had similar body weight to wild type mice at 24 weeks of age that fell within the range considered normal for adult C57/BL6 and 129SvEv mouse lines (Fig. 2A) (37,38).

**Fig. 2.**
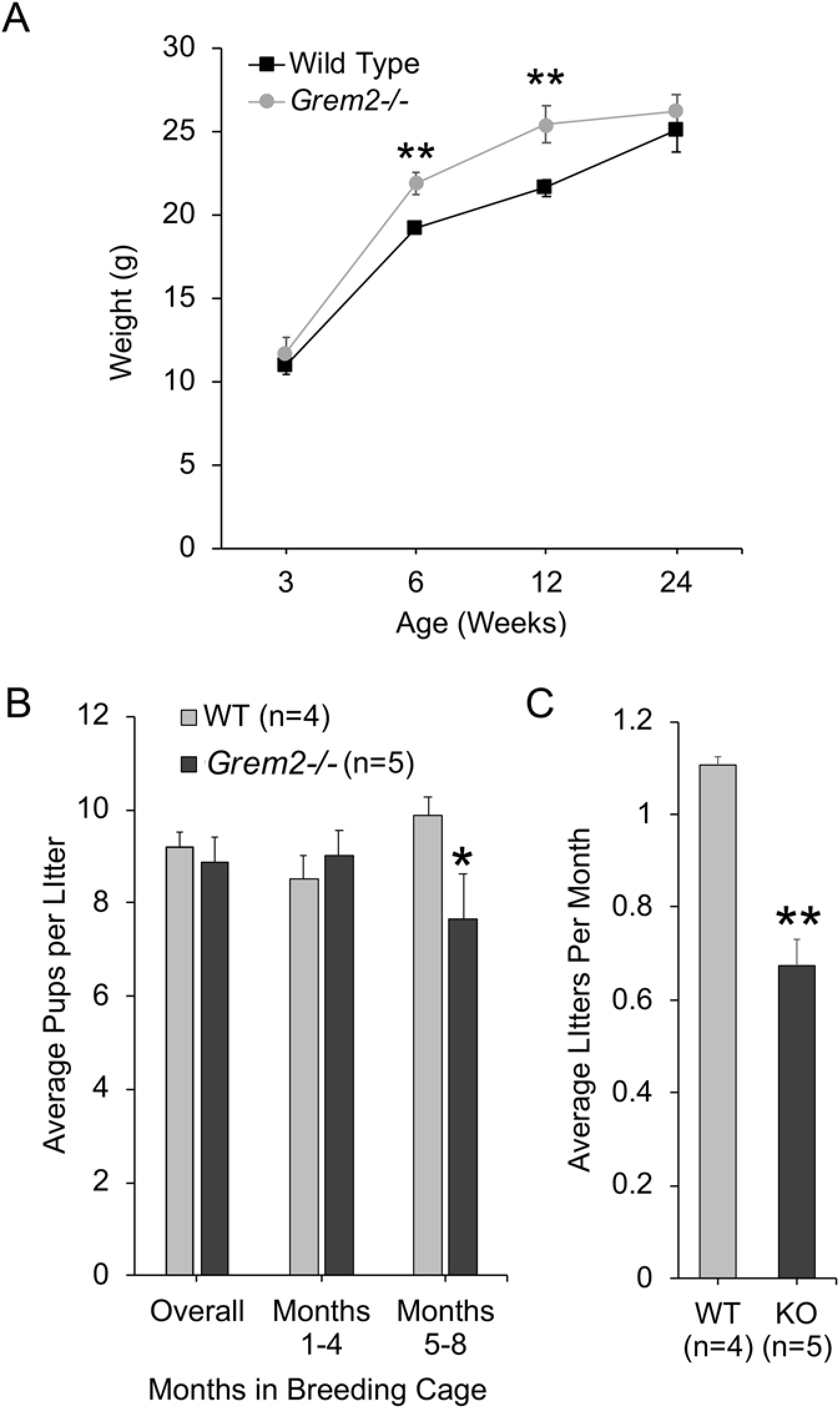
*Grem2^-/-^* females are subfertile. (A) Body weight (g) at time of necropsy for wild type (black squares) and *Grem2^-/-^* (grey circles) females at 3, 6, 12, and 24 weeks of age. Markers represent the mean +/− s.e.m. of n=3-16 animals of each genotype, **P<0.01 by Student’s *t*-test between wild type and knockout mice at the indicated time point. (B) Average litter sizes (pups per litter) wild type (n=4) and *Grem2^-/-^* (n=5) mice. Females of each genotype were set up in breeding pairs at sexual maturity (6-8 weeks of age) to a wild type male and the number of pups recorded over eight months. The average pups per litter is shown as mean +/− s.e.m for the entire breeding trial (“overall”; months 1-8) or in two age brackets (1-4 months versus 5-8 months). The data were split into two age brackets to determine if there was any effect of age, with the asterisk indicated statistical significance in the 5-8-month group; *P<0.05 by Student’s *t-*test. (C) Average litters per month between wild type and *Grem2^-/-^* (“KO”) females in the same 8-month breeding trial as shown in panel B. Data are shown as mean +/− s.e.m, with **indicating P<0.01 by Student’s *t*-test.

### Loss of *Grem2* reduced female fecundity

To test the effect of *Grem2* loss on female fertility, 6-8-week old WT and *Grem2^-/-^* female mice were pair bred to fertile males continuously for eight months and numbers of pups per litter and litters per month were recorded. Overall, wild type females gave birth to an average of 9.2 +/− 0.3 pups per litter and 1.1 +/− 0.02 litters per month (Fig. 2B, C). *Grem2^-/-^* females gave birth to similar numbers of pups per litter (8.9 +/− 0.5). However, if age is considered, a small but statistically significant, decline of 22% in pups per litter was detected in older *Grem2^-/-^* females when the breeding data were split into two age groups (younger and older) (P=0.03) (Fig. 2B). In addition, there was a significant decrease in the numbers of litters per month in *Grem2^-/-^* females (0.67 +/− 0.05) compared to wild type females (1.02 +/− 0.02) (P=0.001) (Fig. 2C). Cycle irregularity in *Grem2^-/-^* occurred regardless of age; on average, during months 1-4, *Grem2^-/-^* female mice missed an average of 1.2 +/− 0.2 litters (compared to 0 for the wild type), and during months 5-8, *Grem2^-/-^* female mice missed 1.8 +/− 0.2 litters (compared to 0 for the wild type). Thus, loss of *Grem2* caused an overall reduction in fecundity that appears to be primarily driven by a reduction in litter production.

### *Grem2^-/-^* females have abnormal estrous cycles

To determine if there were defects in ovarian function, we first analyzed the ovarian histology of wild type and *Grem2^-/-^* females at multiple time points, including immature (3-weeks of age) and sexually mature (6-weeks, 12-weeks, and 6-months of age). In sexually immature 3week old mice, there were no significant differences in the total number of ovarian follicles at the primordial, primary, secondary, or antral stage comparing wild type to *Grem2^-/-^* ovaries (Fig. 3A) and ovaries were of similar size with no obvious histologic defects (Fig. 3C-D). However, there was a statistically significant reduction (P<0.01) in the number of atretic follicles in *Grem2^-/-^* compared to wild type (Fig. 3B). At 6 or 12 weeks of age, there were no gross histologic differences in ovaries from adult *Grem2^-/-^* females compared to wild type mice all stages of follicles including corpora lutea (CL) were present in both genoytypes (data not shown).

**Fig. 3.**
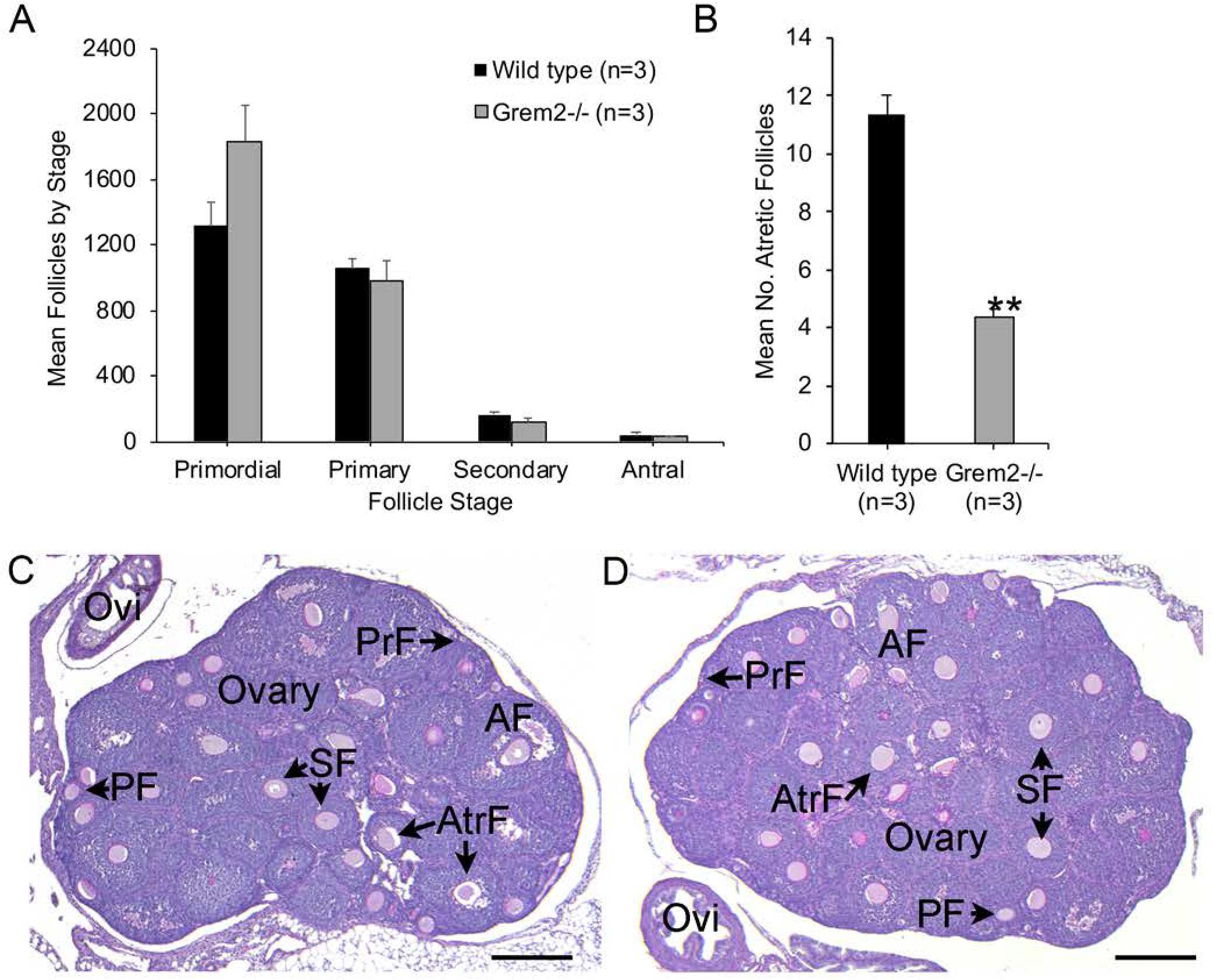
*Grem2^-/-^* ovaries show normal follicle morphometrics prior to sexual maturity. (A) (A) Morphometric assessment of follicle numbers in sexually immature mice. Shown are mean +/− s.e.m. of follicle counts obtained from n=3 of each genotype at 3-weeks of age. B) Atretic follicle counts for the same ovaries as in panel A but at a different scale due to the low overall amounts of follicle atresia in immature mice. **P<0.01 indicates a significantly reduced number of atretic follicles in *Grem2^-/-^* by Student’s *t*-test. (C) Representative PAS histology for a 3-week wild type ovary showing primordial follicles (PrF), primary follicles (PF), secondary follicles (SF) and antral follicles (AF). The oviduct (OVI) is also indicated. (D) Representative PAS histology for a 3-week old Grem2^*-/-*^ with similar follicle stages as the wild type. Scale bar in C, D 100 μm.

At 6-months of age, both wild type and *Grem2^-/-^* ovaries contained follicles of each size class (primordial, primary, secondary, antral) as well as CL (Fig. 4A, B). However, 75% of *Grem2^-/-^* (n=4) contained unusually large patches of PAS positive regions of multinucleated cells, while wild type ovaries (n=4) showed only minimal patches (Fig. 4C, D). These large patches of PAS+ cells have been previously described as multinucleated macrophage giant cells that are typically present in ovaries from aged mice but mostly absent from mice less than 7 months of age (39). By immunohistochemistry, these cells were positive for the mouse macrophage marker F4/80 (Fig. 4E, F). Previous studies have also shown that in aged (*i.e.*, +21 weeks) mouse ovaries, the presence of macrophage giant cells is associated with increased fibrosis and inflammation (39). To identify if *Grem2^-/-^* ovaries showed increased fibrosis, 6-month old ovary sections were stained with picrosirius red (PSR), which has previously been validated as an indicator of fibrous collagen in the ovary (39). A similar level of PSR staining was observed between wild type and *Grem2^-/-^* ovaries (data not shown), indicating similar levels of fibrous collagen in the ovaries, and thus macrophage infiltration does not appear to be directly linked to fibrosis, unlike in aged mice (39). As a prior study of the role of *Grem2^-/-^* in the heart after myocardial infarction (MI) (40), identified increases in inflammatory markers, which included *Bmp2*, tumor necrosis factor alpha (*Tnf*) and e-selectin (*Sele*), we therefore analyzed potential upregulation of these genes in 6-month old ovaries from WT and *Grem2^-/-^* by qPCR; however, we did not see any difference in expression levels (Table 1).

**Fig. 4.**
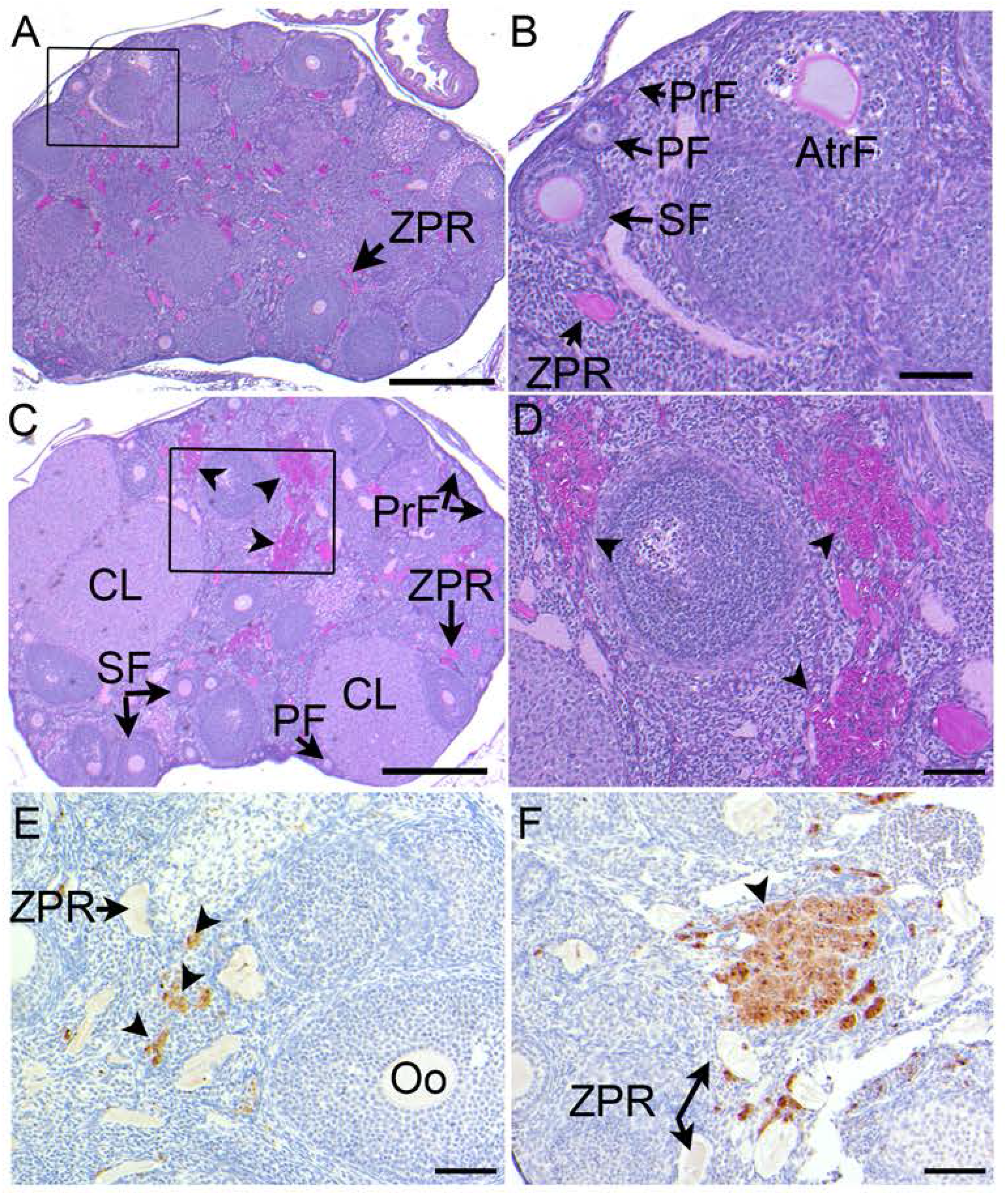
Sexually mature *Grem2^-/-^* ovaries show continuing folliculogenesis but evidence of macrophage infiltration at six months of age. (A, B) Representative PAS histology of a wild type ovary at 6-months of age showing folliculogenesis and oocyte remnants (ZPR) within the interior of the ovary, which is typical in this strain of mice. Area boxed in (A) is shown at a higher magnification in panel B. (B) Boxed area in panel (A) showing primordial follicles (PrF), primary follicles (PF), secondary follicles (SF), ZPR, and an atretic follicle (Atr) are shown at higher magnification for the representative wild type ovary. (C) Representative PAS histologic section of an ovary from a 6-month old *Grem2^-/-^* mouse. All follicle stages are present as are corpora lutea (CL), indicating ovulation. Boxed areas in panel C is shown as higher magnification images in panel D. Arrowheads in panel C, D indicate large patches of PAS+ cells that are distinct from PAS+ ZPRs (Panel E). (E) Anti-F4/80 immunohistochemistry in a representative wild type ovary showing the typical pattern of single positive cells (arrowheads) scattered within the stroma, theca, and CL. (F) Representative *Grem2^-/-^* ovary at the same magnification as panel (E) showing regions of F4/80 positive immunoreactivity (arrowheads) in areas that are larger and less dispersed as well as an F4/80 negative ZPR. Scale bar in panels A, C, 200 μm; panel B, D,E, F, scale bar 50μm.

**Table 1.** Summary of gene expression by qPCR of six-month-old wild type (n=6) and *Grem2^-/-^* (n=6) ovaries at diestrus. Data were analyzed by the ΔΔCT method using *Gapd* for normalization and data are shown relative to the wild type mean. Fold change is levels of *Grem2^-/-^* compared to WT. Data were analyzed using the nonparametric Mann-Whitney *U* test. ^a^Statistical significance between wild type and *Grem2^-/-^* ovaries (P<0.05). ND, no difference.

| Symbol | Gene | WT<br>Ave. +/-<br>s.e.m. | *Grem2*<sup>-/-</sup><br>Ave. +/- s.e.m. | Fold<br>Change |
| --- | --- | --- | --- | --- |
| *Bmp2* | Bone morphogenetic protein-2 | 1.0 +/- 0.2 | 1.0 +/- 0.1 | ND |
| *Bmp4* | Bone morphogenetic protein-4 | 1.0 +/- 0.3 | 0.7 +/- 0.1 | ND |
| *Bmp15* | Bone morphogenetic protein-15 | 1.0 +/- 0.2 | 1.2 +/- 0.1 | ND |
| *Cyp19a1* | Aromatase | 1.0 +/- 0.3 | 0.8 +/- 0.1 | ND |
| *Fshr* | Follicle stimulating hormone receptor | 1.0 +/- 0.3 | 1.0 +/- 0.0 | ND |
| *Gdf9* | Growth and differentiation factor-9 | 1.0 +/- 0.1 | 1.1 +/- 0.1 | ND |
| *Kitl1* | Kit ligand 1 | 1.0 +/- 0.1 | 0.8 +/- 0.1 | ND |
| *Kitl2* | Kit ligand 2 | 1.1 +/- 0.1 | 0.9 +/- 0.2 | ND |
| *Sele* | Selectin E | 1.0 +/- 0.2 | 1.0 +/- 0.1 | ND |
| *Tnfa* | Tumor necrosis factor $\alpha$ | 1.0 +/- 0.2 | 0.9 +/- 0.1 | ND |

Because of the significant changes to litter production, (Fig. 2C), estrous cycles were evaluated in six-month-old females. Six-month old wild type mice had normal estrous cycles that averaged 4-5 days while *Grem2^-/-^* female mice had irregular estrous cycles (Fig. 5A). *Grem2^-/-^* females had a significant increase in the percentage of days spent in metestrus and diestrus and a concomitant decrease in the number of estrous cycles per month (Fig. 5B, C). Serum hormone profiles of diestrous stage 6-month old females showed no change in mean levels of estradiol, testosterone, luteinizing hormone, or follicle stimulating hormone (FSH) between wild type and *Grem2^-/-^* (Table 2). We additionally used linear regression to test the correlation between estradiol and FSH levels. While diestrous stage wild type mice exhibit a correlation between estradiol and FSH (r^2^=0.48), this correlation is not found in diestrous stage *Grem2^-/-^* females (r^2^=0.07) (Fig. 6A).

**Fig. 5.**
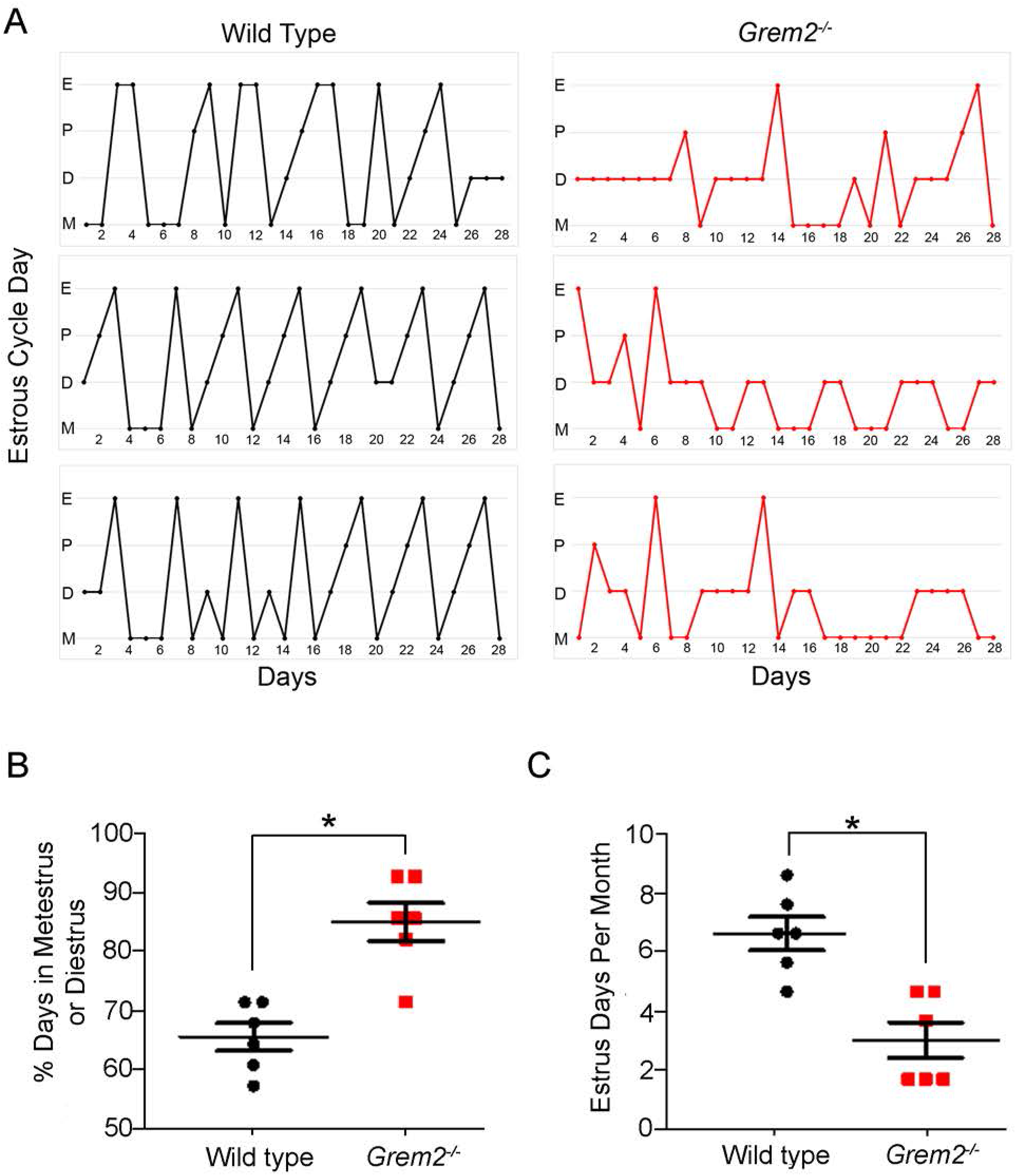
*Grem2^-/-^* mice have irregular estrous cycles. (A) Three typical estrous cycles for each genotype are shown for 6-month old wild type (WT) (black) and *Grem2^-/-^* (red) mice from a total of n=6 mice per genotype. Estrous cycle day is indicated on the y-axis as estrus (E), proestrus (P), diestrus (D), and metestrus (M) for both genotypes (B) Percentage of time in metestrus and diestrus, and (C) number of days in estrus for 6-month old wild type (n=6) and *Grem2^-/-^* (n=6) for a one-month period. *P<0.05.

**Fig. 6.**
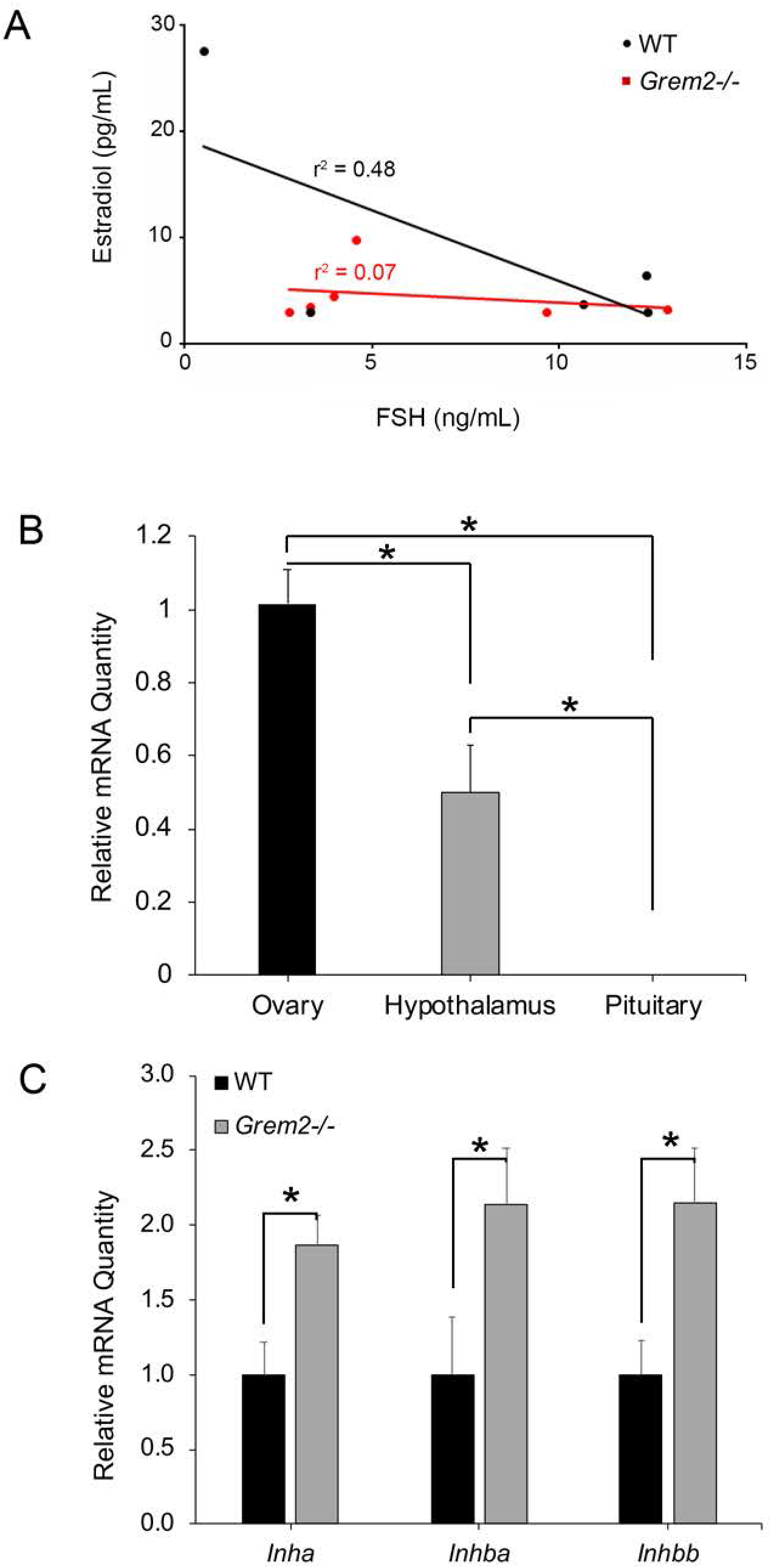
Lack of correlation between serum estradiol and FSH in diestrous stage *Grem2^-/-^* mice. (A) Linear regression of serum estradiol (pg/mL) and FSH (ng/mL) in WT (shown as black circles) (n=5) and *Grem2^-/-^* (shown as red squares) (n=6) at six months of age. (B) Relative mRNA expression levels by qPCR for *Grem2* in the wild type mouse ovary, hypothalamus, and pituitary normalized to *Gapd* and relative to the level of *Grem2* in the ovary (n=5 animals). (C) mRNA expression of inhibin/activin subunits, *Inha, Inhba, Inhbb*, by qPCR in ovaries from diestrus stage 6-month-old mice (n=6 each genotype). *P<0.05 by Student’s *t*-test.

**Table 2.** Serum hormone data for six-month old female wild type and *Grem2^-/-^* mice at diestrus. Results are shown as the mean +/− s.e.m. for the indicated number of females (n). Data were log transformed prior to statistical analysis using the nonparametric Mann-Whitney *U* test. Statistical significance was only detected for AMH values, **P<0.01

| Hormone | Wild type (n) | *Grem2*<sup>-/-</sup> (n) |
| --- | --- | --- |
| FSH (ng/mL) | 7.8 +/- 2.5 (5) | 6.2 +/- 1.7 (6) |
| LH (ng/mL) | 0.4 +/- 0.1 (5) | 1.1 +/- 0.8 (6) |
| Estradiol (pg/mL) | 8.7 +/- 4.8 (5) | 4.5 +/- 1.1 (6) |
| Testosterone (ng/dL) | 45.5 +/- 29.8 (5) | 31.0 +/- 5.6 (6) |
| Anti-Müllerian Hormone (ng/mL) | 146.8 +/- 13.8 (5) | 66.1 +/- 10.5** (6) |

Alterations in the estrous cycle could be due to dysfucntion of the HPO axis, but there is limited information for a role for *Grem2*, even though BMP4 is known to be important in mouse pituitary development (41) and may regulate FSH in gonadotropes (42,43). *Grem2* is expressed in the brain (1), but hypothalamic expression has not been previously reported (44). By qPCR, *Grem2* could not be detected in the adult mouse pituitary; however, expression was present in the hypothalamus at ~60% relative to the ovary (Fig. 6B). As the production of the peptide hormone, inhibin, from the ovary is also responsible for HPO negative feedback and suppression of FSH, ovarian expression of genes encoding the inhibin/activin subunits were measured. By qPCR, transcript levels of *Inha, Inhba*, and *Inhbb* significantly increased (P<0.05) in *Grem2^-/-^* ovaries compared to wild type (Fig. 6C). Markers of key cell types in the ovary showed no difference between genotypes, including genes expressed in oocytes (*Bmp15, Gdf9*), granulosa cells (*Bmp2, Cype19a1, Fshr, Kitl1/2*) or thecal cells (*Bmp4*) (Table 1), supporting the histologic findings of similar follicle stages at this age. Because of its role as a marker of the ovarian reserve (45–47), serum levels of AMH were measured in wild type and *Grem2^-/-^* mice.

Compared to wild type, *Grem2^-/-^* showed significantly reduced levels of AMH at 6 months of age (P<0.01) (Table 2). As AMH is secreted from granulosa cells of growing follicles, AMH immunoreactivity in individual follicles was quantified by immunofluorescence (Fig. 7). When plotted by mean fluorescent intensity versus follicle stage, there was significantly reduced AMH immunoreactivity in *Grem2^-/-^* preantral follicles mice compared to the follicles of wild type mice (Fig. 7), indicating that it is not loss of follicles that reduces serum AMH, but instead, the reduction is due to decreased expression in granulosa cells.

**Fig. 7.**
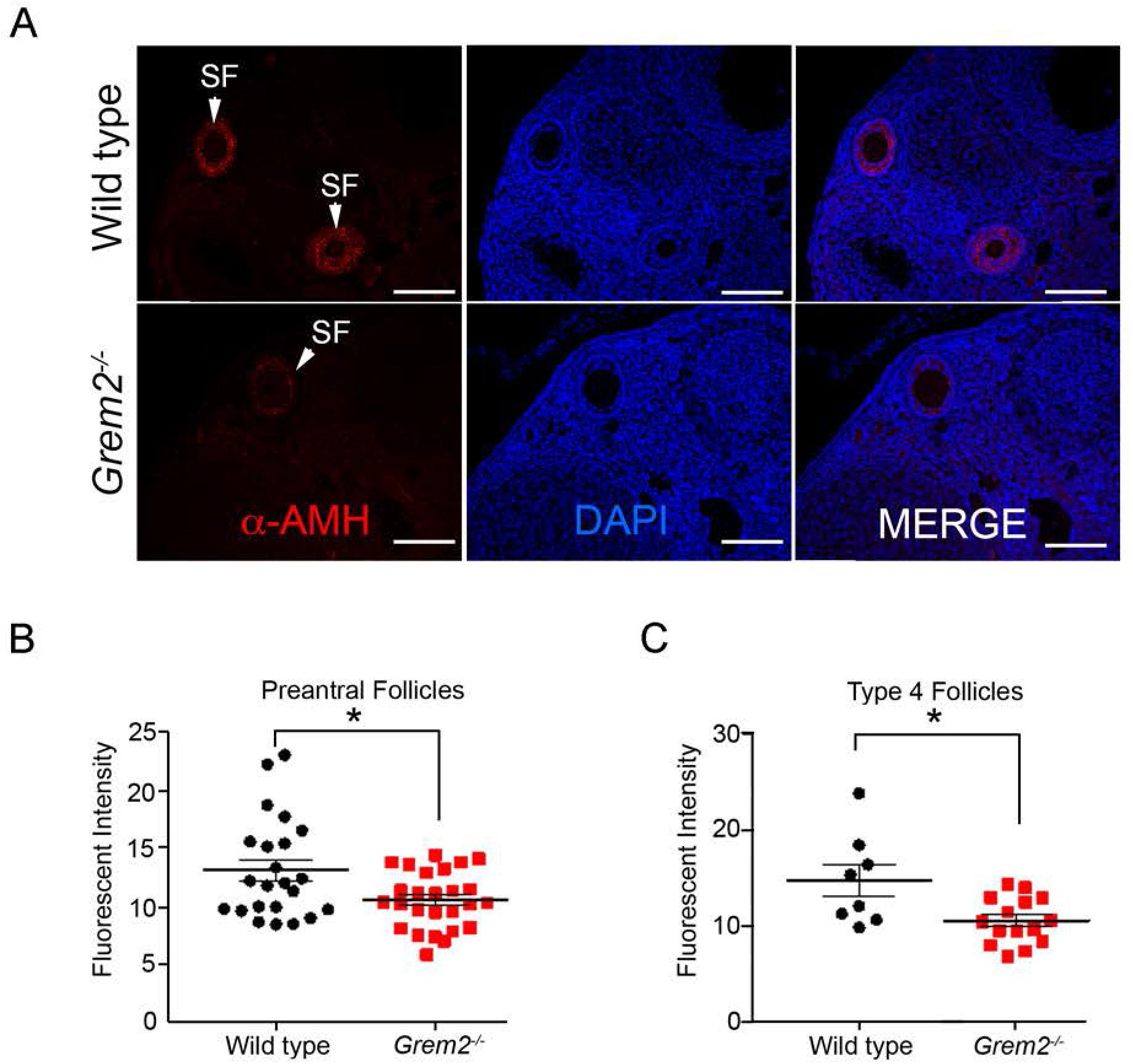
Follicles from *Grem2^-/-^* mice produce reduced levels of AMH. Ovary sections from 6- month-old WT and *Grem2^-/-^* mice were analyzed for AMH immunoreactivity (red) and DAPI (blue) was used to stain nuclei (A). Arrowheads indicate secondary follicles (SF). Scale bars, 100 μm (n=3 per genotype). Data were analyzed by classifying follicles as preantral follicle type (B) or by individual types (C, D, and E). Grouping all preantral follicles showed lower levels of expression in the *Grem2^-/-^* mice (B), as did type 4 follicles (D).

### *GREM2* has a rare variant in patient samples of POI

A previously published cohort of POI patient samples that had undergone whole exome sequencing (WES) was queried for nonsynonymous and splice site variants in GREM2 (30). Of the 103 POI cases sequenced, one individual contained a single novel nonsynonymous heterozygous variant in *GREM2*, c.C356T:p.S119F in exon 2. This variant was not present in The Genome Aggregation Database (gnomAD) or other databases. We further modeled the location of this variant based on the previously published crystal structure of GREM2 with GDF5 (6). The S119F variant lies in the interface of the interaction domain between antagonist and ligand (Fig. 8), which has previously been shown to be key region required for robust BMP antagonism (6).

**Fig. 8.**
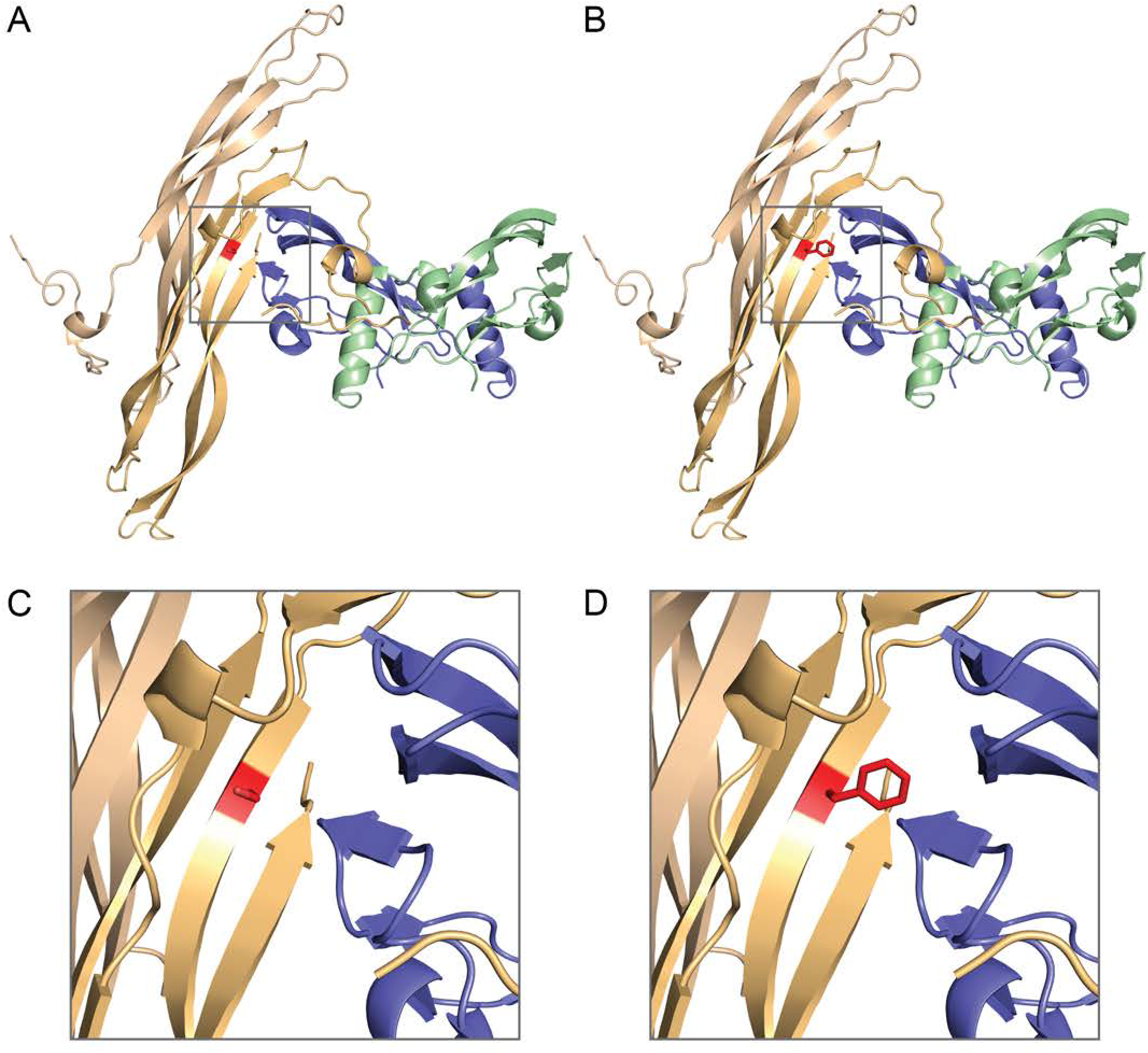
Model of the GREM2 variant S119F with the ligand GDF5. (A) Structure of GREM2 (monomers in pale orange and tan) in complex with GDF5 (monomers in slate and pale green) with S119 shown in red, PDB ID: 5HK5 (6) (B) Structure of GREM2-GDF5 with S119 mutated to F119, with the most probable rotomer shown in red. Zoomed in view of panel A, focusing on the S119 residue and its local interactions. (D) Zoomed in view of panel B, focusing the F119 residue and its local interactions.

## Discussion

Because of their roles as powerful developmental morphogens and regulators of adult tissue homeostasis, BMP activity is under strict biologic control. One mechanism for their regulation is through production of extracellular binding proteins, including GREM2, which when bound to BMPs, disrupts the ability of the ligand to form the ternary signaling receptor complex. GREM2 exists as a stable non-disulfide bonded dimer with binding affinities for BMP2 and BMP4 in the nanomolar range (1,2,6,48). Additionally, GREM2 binds and inhibits AMH, another member of the TGFβ family, in *in vitro* assays (7). *GREM2* is expressed in the fetal human ovary as well as in granulosa cells of mouse ovarian follicles (1,16) although its role in either the embryonic or adult ovary is not well understood. A previous mouse knockout of *Grem2* was published as part of a high-throughput knockout phenotyping program by Lexicon Pharmaceuticals that included fertility assays. These assays typically were performed from ages 8-16 weeks using two homozygous knockout females mated to a wild type male (49). The major defect identified in Lexicon’s *Grem2^-/-^* line related to small and malformed upper and lower mandibular incisors (14). No fertility defects were noted, though it is unlikely that the fertility screen had sufficient depth to identify changes in fecundity beyond overt sterility, particularly for those that arise due to aging. As that model was unavailable, we developed a new *Grem2^-/-^* mouse model using CRISPR/Cas9 gene targeting to delete 1kb containing the entire coding exon (exon 2). This new mouse model phenocopies the Lexicon deletion with respect to dental defects, even though the genetic background is dissimilar [inbred C57/Bl6 (albino) versus mixed hybrid in our study], suggesting a robust phenotype resulting from loss of *Grem2* in tooth development.

Unlike homozygous mutations in *Grem1*, which are perinatal lethal (27,50), *Grem2^-/-^* mice are viable but subfertile. Overall litter production in *Grem2^-/-^* females was reduced throughout the reproductive lifespan, but appears to worsen in older mice (i.e., 6-8 months of age). This change in litter production primarily results from irregular estrous cycles. As *Grem2* is widely expressed and transcripts have been identified in the mouse ovary, brain, and uterus amongst other organs (1), it is currently unclear if the changes in cyclicity are due intraovarian defects, defects in other tissues, or a combination of both. Within the mouse ovary, *Grem2* is expressed from granulosa cells and is upregulated in response to gonadotropin stimulation (1,27,51). While *Grem2* expression could not be detected in the mouse pituitary by qPCR, mRNA transcripts were detected in the hypothalamus at about half the level in the ovary. The relative contribution of intraovarian defects versus potential hypothalamic defects remains to be determined and would require generation of a conditional allele for *Grem2* for cell-specific deletion. Interestingly, the AMH receptor (*Amhr*) has been detected in a subset of gonadotropin releasing hormone (GnRH) neurons within the hypothalamus and at least one study has shown that AMH potently activates GnRH neuron firing and GnRH-dependent LH pulsatility and secretion (52). If GREM2 regulates AMH activity as has been suggested (7), then it is possible that GnRH neuronal activity may be disrupted in *Grem2^-/-^* females and contribute to the fertility defect. Such studies require more precise measurements of LH pulse generation, rather than the steady state (diestrus) levels reported here.

Previous studies suggest GREM2 has a role in embryonic human ovary development, as its expression increases between 8-11 weeks and 14-16 weeks gestation, which corresponds to the timing of post-migratory germ cell proliferation and entry into meiosis I, respectively (16). This study also demonstrates that GREM2 partially antagonizes BMP4 induced gene expression (16). Furthermore, treatment of organ cultures of rat ovaries with GREM2 reverses the ability of AMH to suppress primordial follicle activation (7). Surprisingly, ovaries from sexually immature (3-week old) *Grem2^-/-^* mice contain equivalent numbers of primordial follicles as the wild type mice, suggesting that *Grem2* may not play a major role in embryonic or postnatal formation of the ovarian reserve in mice, or that developmental changes in the ovarian reserve are resolved to wild type levels during the first wave of folliculogenesis. This is different than what has been noted for mice homozygous null for *Grem1*, which have decreased numbers of germ cells and primordial follicles at birth (27). Alternatively, loss of *Grem2* in the embryo or postnatally could be compensated for by another BMP antagonist, such as *Grem1*. Functional redundancy between *Grem2* and *Grem1* has been hypothesized (7,27), but not demonstrated.

Ovaries from adult *Grem2^-/-^* contained all stages of follicle growth, including primordial follicles, and produced normal-sized litters earlier in their reproductive lifespan but show a small but significant decrease in litter sizes (~24%) at later ages (>6 months of age). This suggests some intraovarian defect in older mice. While the mean diestrous stage serum levels of estradiol and FSH were similar between wild type and *Grem2^-/-^* mice, the well-known correlation between estradiol and FSH was altered in *Grem2^-/-^* females. In wild type rodents, at metestrus and diestrus, low but rising levels of estradiol from granulosa cells of growing follicles negatively regulate production of FSH by suppressing hypothalamic secretion of gonadotropin hormone releasing hormone (53–55). Furthermore, circulating levels of inhibin suppress FSH production from the pituitary (55–57). In *Grem2^-/-^* ovaries, the inhibin/activin subunit genes (*Inha, Inhba, Inhbb*) were upregulated in adult animals, which may further contribute to changes to reproductive cyclicity if circulating levels of inhibin A and inhibin B, which are known to vary with the estrous cycle stage in rodents (57), are also altered. A more detailed analysis of FSH and LH production and secretion as well as an analysis of circulating inhibin levels during the different stages of the estrous cycle will help to resolve these issues. In addition, in *Grem2^-/-^* ovaries, there were loss of granulosa cell production of AMH, which is most highly expressed in growing preantral follicles (58,59). Suppression of *Amh* expression in *Grem2^-/-^* ovaries could be directly or indirectly related to changes in BMP signaling, as regulation of *Amh* by various growth factor pathways during preantral folliculogenesis is not fully understood (60). *Amh* is known to be downregulated in rodents and human when follicles reach the antral stage (61), so potentially, the *Grem2^-/-^* model will allow us to elucidating key pathways controlling AMH production.

While litter production was reduced in *Grem2^-/-^* females after 10-12 weeks of age compared to wild type mice, it was more pronounced after six months of age. However, unlike wild type mice, 6-month old *Grem2^-/-^* ovaries accumulate large patches of F4/80+ multinucleated macrophage giant cells, which were not present at three-month of age (data not shown). The significance of macrophage giant cells is unknown, but ovaries from aged wild type female mice (14-17 months of age) accumulate areas of fibrotic tissue concurrent with chronic inflammation and macrophage giant cells and these were generally absent in ovaries from mice less than 7 months of age (CD1 and CB6F1 strains) (62). Thus *Grem2^-/-^* ovaries have one hallmark of early aging, but without significant changes in fibrosis. The presence of large patches of F4/80+ macrophages also suggests an increase in tissue inflammation, similar to a previous study of *Grem2^-/-^* within the heart the context of recovery from MI. Following MI, *Grem2^-/-^* mice show excessive inflammation, including increases in F4/80+ macrophages and poorer functional outcomes as result of overactive BMP signaling, which can be rescued by intraperitoneal administration of recombinant GREM2 (40). Furthermore, a BMP pro-inflammatory cascade has been suggested in other diseases, including chronic inflammatory arthritis and atherosclerosis, though in other diseases BMP signaling is anti-inflammatory (63,64). How loss of *Grem2* affects the balance of TGFβ superfamily signaling, including AMH and BMPs, within the ovary remains to be determined. As AMH has reduced expression in follicles of *Grem2^-/-^* ovaries, inter-and intra-follicle BMP signaling may predominate over AMH, promoting macrophage recruitment or differentiation, disrupting ovarian function, and possibly altering feedback within the HPO axis.

A number of recent studies implicate changes to BMP signaling with the development of POI in women. This includes genetic variants in BMP ligands (*BMP15*) (65,66), the BMP15 promoter (67), the BMP receptors (*BMPR1A* and *BMPR1B*) (68), and the BMP antagonist, *GREM1* (15). It is currently unknown how these variants contribute to POI in women. Because these genes are also expressed in the brain and pituitary in various species (42,69–72), alterations in their activity in the etiology of POI could occur at multiple levels of the HPO axis. Studies on the structure of GREM2 with GDF5 indicates that GREM2 forms alternating higher form stable aggregates with its ligands that is unique among the BMP antagonists, as compared to Noggin and Follistatin (6). The location of the S119F mutation lies within the interface between GREM2 and GDF5, but it is currently not known how this mutation affects functional antagonism. Mechanistically, GREM2 wraps around the BMP signaling molecule, occluding both the type I (located at the concave dimer interface) and type II (at the convex surface of the ligand) binding sites needed for signaling (6). The side chain of the S119 residue points directly into one of the primary binding interfaces of GDF5, specifically the one that impairs binding of GDF5 to the type II BMP receptors. Previous mutational work has demonstrated that this section of GREM2 is particularly important to robust BMP inhibition and sensitive to mutational disruption (6). Additionally, when we modeled S119 with a phenylalanine, a significant steric clash occurred with both the ligand and residues of GREM2 important for GDF5 binding. Thus, we anticipate the S119F mutation would interfere ligand binding and weaken GREM2 antagonism, potentially leading to a gain-of-function in BMP or AMH signaling. The genetic variants found for *BMPR1A (ALK3*) and *BMPR1B (ALK6*) are located within the kinase domain of the receptors, and show altered signaling when measured by *in vitro* assays. The BMPR1A p.Arg442His variant inhibits receptor activation and was discovered in a patient with menarche at 14 years and amenorrhea 1 year later (73). The BMPR1B p.Phe272Leu variant shows constitutive activation when tested *in vitro*, and was discovered in a patient with secondary amenorrhea at age 27 (73). Thus, both loss-of-function and gain-of-function mutations that alter BMP signaling may drive POI. Of interest, mouse knockouts do not always fully phenocopy human disease; for instance, women with loss-of-function variants in *DCAF17* show hypogonadism, but when the same mutations are made in mouse models, the mice are subfertile (74). Additional structure-function studies will be required to understand how the *GREM2* variant alters BMP or AMH signaling and its consequences on HPO function.

## Acknowledgements

Core services at Baylor College of Medicine are supported by funding from the NIH (P30 CA125123) and the authors additionally thank members of the core facilities at BCM for their assistance with these studies: Genetically Engineered Mouse Core, the Human Tissue Acquisition and Pathology Core, Embryonic Stem Cell Core and Dr. Jason Heaney), the Vital Microscopy and in *vivo* Imaging Core, and the Integrated Microscopy Core. The authors additionally acknowledge the University of Virginia Center for Reproductive Research Ligand Assay and Analysis Core (supported by NIH R24 HD102061) for hormone analysis. We thank Ramya Masand, MD (Baylor College of Medicine) for histologic assessment of ovaries and Shailaja K. Mani, PhD (Baylor College of Medicine) for assistance with dissection of mouse hypothalami.

## References

1. Sudo S, Avsian-Kretchmer O, Wang LS, Hsueh AJ. Protein related to DAN and cerberus is a bone morphogenetic protein antagonist that participates in ovarian paracrine regulation. J Biol Chem. 2004;279(22):23134–23141.

2. Nolan K, Kattamuri C, Luedeke DM, Deng X, Jagpal A, Zhang F, Linhardt RJ, Kenny AP, Zorn AM, Thompson TB. Structure of protein related to Dan and Cerberus: insights into the mechanism of bone morphogenetic protein antagonism. Structure. 2013;21(8):1417–1429.

3. Kattamuri C, Luedeke DM, Thompson TB. Expression and purification of recombinant protein related to DAN and cerberus (PRDC). Protein Expr Purif. 2012;82(2):389–395.

4. Minabe-Saegusa C, Saegusa H, Tsukahara M, Noguchi S. Sequence and expression of a novel mouse gene PRDC (protein related to DAN and cerberus) identified by a gene trap approach. Dev Growth Differ. 1998;40(3):343–353.

5. Gazzerro E, Canalis E. Bone morphogenetic proteins and their antagonists. Rev Endocr Metab Disord. 2006;7(1-2):51–65.

6. Nolan K, Kattamuri C, Rankin SA, Read RJ, Zorn AM, Thompson TB. Structure of Gremlin-2 in Complex with GDF5 Gives Insight into DAN-Family-Mediated BMP Antagonism. Cell Rep. 2016;16(8):2077–2086.

7. Nilsson EE, Larsen G, Skinner MK. Roles of Gremlin 1 and Gremlin 2 in regulating ovarian primordial to primary follicle transition. Reproduction. 2014;147(6):865–874.

8. Kattamuri C, Nolan K, Thompson TB. Analysis and identification of the Grem2 heparin/heparan sulfate-binding motif. Biochem J. 2017;474(7):1093–1107.

9. Lu L, Huang J, Xu F, Xiao Z, Wang J, Zhang B, David NV, Arends D, Gu W, Ackert-Bicknell CL, Sabik OL, Farber CR, Quarles LD, Williams RW. Genetic dissection of femoral and tibial microarchitecture. JBMR Plus. 2019;3(12):e10241.

10. Kaminski A, Bogacz A, Uzar I, Czerny B. Association betweenGREM2 gene polymorphism with osteoporosis and osteopenia in postmenopausal women. Eur J Obstet Gynecol Reprod Biol. 2018;228:238–242.

11. Muller, II, Melville DB, Tanwar V, Rybski WM, Mukherjee A, Shoemaker MB, Wang WD, Schoenhard JA, Roden DM, Darbar D, Knapik EW, Hatzopoulos AK. Functional modeling in zebrafish demonstrates that the atrial-fibrillation-associated gene GREM2 regulates cardiac laterality, cardiomyocyte differentiation and atrial rhythm. Dis Model Mech. 2013;6(2):332–341.

12. Mostowska A, Biedziak B, Zadurska M, Bogdanowicz A, Olszewska A, Cieslinska K, Firlej E, Jagodzinski PP. GREM2 nucleotide variants and the risk of tooth agenesis. Oral Dis. 2018;24(4):591–599.

13. Cheung CL, Lau KS, Sham PC, Tan KC, Kung AW. Genetic variants in GREM2 are associated with bone mineral density in a southern Chinese population. J Clin Endocrinol Metab. 2013;98(9):E1557–1561.

14. Vogel P, Liu J, Platt KA, Read RW, Thiel M, Vance RB, Brommage R. Malformation of Incisor Teeth in Grem2-/-Mice. Vet Pathol. 2014.

15. Patino LC, Beau I, Carlosama C, Buitrago JC, Gonzalez R, Suarez CF, Patarroyo MA, Delemer B, Young J, Binart N, Laissue P. New mutations in non-syndromic primary ovarian insufficiency patients identified via whole-exome sequencing. Hum Reprod. 2017:1–9.

16. Bayne RA, Donnachie DJ, Kinnell HL, Childs AJ, Anderson RA. BMP signalling in human fetal ovary somatic cells is modulated in a gene-specific fashion by GREM1 and GREM2. Mol Hum Reprod. 2016;22(9):622–633.

17. Nilsson EE, Skinner MK. Bone morphogenetic protein-4 acts as an ovarian follicle survival factor and promotes primordial follicle development. Biol Reprod. 2003;69(4):1265–1272.

18. Durlinger AL, Gruijters MJ, Kramer P, Karels B, Ingraham HA, Nachtigal MW, Uilenbroek JT, Grootegoed JA, Themmen AP. Anti-Mullerian hormone inhibits initiation of primordial follicle growth in the mouse ovary. Endocrinology. 2002;143(3):1076–1084.

19. Durlinger AL, Kramer P, Karels B, de Jong FH, Uilenbroek JT, Grootegoed JA, Themmen AP. Control of primordial follicle recruitment by anti-Mullerian hormone in the mouse ovary. Endocrinology. 1999;140(12):5789–5796.

20. Myers M, Mansouri-Attia N, James R, Peng J, Pangas SA. GDF9 modulates the reproductive and tumor phenotype of female inha-null mice. Biol Reprod. 2013;88(4):86.

21. Ran FA, Hsu PD, Wright J, Agarwala V, Scott DA, Zhang F. Genome engineering using the CRISPR-Cas9 system. Nat Protoc. 2013;8(11):2281–2308.

22. Bassett AR, Tibbit C, Ponting CP, Liu JL. Highly efficient targeted mutagenesis of Drosophila with the CRISPR/Cas9 system. Cell Rep. 2013;4(1):220–228.

23. Byers SL, Wiles MV, Dunn SL, Taft RA. Mouse estrous cycle identification tool and images. PLoS One. 2012;7(4):e35538.

24. Pangas SA, Jorgez CJ, Tran M, Agno J, Li X, Brown CW, Kumar TR, Matzuk MM. Intraovarian activins are required for female fertility. Mol Endocrinol. 2007;21(10):2458–2471.

25. Tilly JL. Ovarian follicle counts--not as simple as 1, 2, 3. Reprod Biol Endocrinol. 2003;1:11.

26. Whitcomb BW, Schisterman EF. Assays with lower detection limits: implications for epidemiological investigations. Paediatr Perinat Epidemiol. 2008;22(6):597–602.

27. Myers M, Tripurani SK, Middlebrook B, Economides AN, Canalis E, Pangas SA. Loss of gremlin delays primordial follicle assembly but does not affect female fertility in mice. Biol Reprod. 2011;85(6):1175–1182.

28. Pedersen T, Peters H. Proposal for a classification of oocytes and follicles in the mouse ovary. J Reprod Fertil. 1968;17:555–557.

29. Livak KJ, Schmittgen TD. Analysis of relative gene expression data using real-time quantitative PCR and the 2(-Delta Delta C(T)) Method. Methods. 2001;25(4):402–408.

30. Yang X, Touraine P, Desai S, Humphreys G, Jiang H, Yatsenko A, Rajkovic A. Gene variants identified by whole-exome sequencing in 33 French women with premature ovarian insufficiency. J Assist Reprod Genet. 2019;36(1):39–45.

31. Desai S, Wood-Trageser M, Matic J, Chipkin J, Jiang H, Bachelot A, Dulon J, Sala C, Barbieri C, Cocca M, Toniolo D, Touraine P, Witchel S, Rajkovic A. MCM8 and MCM9 Nucleotide Variants in Women With Primary Ovarian Insufficiency. J Clin Endocrinol Metab. 2017;102(2):576–582.

32. Richards S, Aziz N, Bale S, Bick D, Das S, Gastier-Foster J, Grody WW, Hegde M, Lyon E, Spector E, Voelkerding K, Rehm HL, Committee ALQA. Standards and guidelines for the interpretation of sequence variants: a joint consensus recommendation of the American College of Medical Genetics and Genomics and the Association for Molecular Pathology. Genet Med. 2015;17(5):405–424.

33. Genomes Project C, Auton A, Brooks LD, Durbin RM, Garrison EP, Kang HM, Korbel JO, Marchini JL, McCarthy S, McVean GA, Abecasis GR. A global reference for human genetic variation. Nature. 2015;526(7571):68–74.

34. Lek M, Karczewski KJ, Minikel EV, Samocha KE, Banks E, Fennell T, O’Donnell-Luria AH, Ware JS, Hill AJ, Cummings BB, Tukiainen T, Birnbaum DP, Kosmicki JA, Duncan LE, Estrada K, Zhao F, Zou J, Pierce-Hoffman E, Berghout J, Cooper DN, Deflaux N, DePristo M, Do R, Flannick J, Fromer M, Gauthier L, Goldstein J, Gupta N, Howrigan D, Kiezun A, Kurki MI, Moonshine AL, Natarajan P, Orozco L, Peloso GM, Poplin R, Rivas MA, Ruano-Rubio V, Rose SA, Ruderfer DM, Shakir K, Stenson PD, Stevens C, Thomas BP, Tiao G, Tusie-Luna MT, Weisburd B, Won HH, Yu D, Altshuler DM, Ardissino D, Boehnke M, Danesh J, Donnelly S, Elosua R, Florez JC, Gabriel SB, Getz G, Glatt SJ, Hultman CM, Kathiresan S, Laakso M, McCarroll S, McCarthy MI, McGovern D, McPherson R, Neale BM, Palotie A, Purcell SM, Saleheen D, Scharf JM, Sklar P, Sullivan PF, Tuomilehto J, Tsuang MT, Watkins HC, Wilson JG, Daly MJ, MacArthur DG, Exome Aggregation C. Analysis of protein-coding genetic variation in 60,706 humans. Nature. 2016;536(7616):285–291.

35. Sherry ST, Ward MH, Kholodov M, Baker J, Phan L, Smigielski EM, Sirotkin K. dbSNP: the NCBI database of genetic variation. Nucleic Acids Res. 2001;29(1):308–311.

36. Pangas SA, Li X, Umans L, Zwijsen A, Huylebroeck D, Gutierrez C, Wang D, Martin JF, Jamin SP, Behringer RR, Robertson EJ, Matzuk MM. Conditional deletion of Smad1 and Smad5 in somatic cells of male and female gonads leads to metastatic tumor development in mice. Mol Cell Biol. 2008;28(1):248–257.

37. Reed DR, Bachmanov AA, Tordoff MG. Forty mouse strain survey of body composition. Physiol Behav. 2007;91(5):593–600.

38. Laboratory TJ. Body weight information for aged C57/BL/6J 2020.

39. RT R, SM B, H T-S, Pangas S. Gremlin-2 is required for female fertility. submitted.

40. Sanders LN, Schoenhard JA, Saleh MA, Mukherjee A, Ryzhov S, McMaster WG, Jr., Nolan K, Gumina RJ, Thompson TB, Magnuson MA, Harrison DG, Hatzopoulos AK. BMP Antagonist Gremlin 2 Limits Inflammation After Myocardial Infarction. Circ Res. 2016;119(3):434–449.

41. Takuma N, Sheng HZ, Furuta Y, Ward JM, Sharma K, Hogan BL, Pfaff SL, Westphal H, Kimura S, Mahon KA. Formation of Rathke’s pouch requires dual induction from the diencephalon. Development. 1998;125(23):4835–4840.

42. Otsuka F, Shimasaki S. A novel function of bone morphogenetic protein-15 in the pituitary: selective synthesis and secretion of FSH by gonadotropes. Endocrinology. 2002;143(12):4938–4941.

43. Huang HJ, Wu JC, Su P, Zhirnov O, Miller WL. A novel role for bone morphogenetic proteins in the synthesis of follicle-stimulating hormone. Endocrinology. 2001;142(6):2275–2283.

44. Fredolini C, Bystrom S, Pin E, Edfors F, Tamburro D, Iglesias MJ, Haggmark A, Hong MG, Uhlen M, Nilsson P, Schwenk JM. Immunocapture strategies in translational proteomics. Expert Rev Proteomics. 2016;13(1):83–98.

45. Anderson RA, Nelson SM, Wallace WH. Measuring anti-Mullerian hormone for the assessment of ovarian reserve: when and for whom is it indicated? Maturitas. 2012;71(1):28–33.

46. Broer SL, Broekmans FJ, Laven JS, Fauser BC. Anti-Mullerian hormone: ovarian reserve testing and its potential clinical implications. Hum Reprod Update. 2014;20(5):688–701.

47. Visser JA, Schipper I, Laven JS, Themmen AP. Anti-Mullerian hormone: an ovarian reserve marker in primary ovarian insufficiency. Nat Rev Endocrinol. 2012;8(6):331–341.

48. Nolan K, Kattamuri C, Luedeke DM, Angerman EB, Rankin SA, Stevens ML, Zorn AM, Thompson TB. Structure of neuroblastoma suppressor of tumorigenicity 1 (NBL1): insights for the functional variability across bone morphogenetic protein (BMP) antagonists. J Biol Chem. 2015;290(8):4759–4771.

49. Brommage R, Liu J, Hansen GM, Kirkpatrick LL, Potter DG, Sands AT, Zambrowicz B, Powell DR, Vogel P. High-throughput screening of mouse gene knockouts identifies established and novel skeletal phenotypes. Bone Res. 2014;2:14034.

50. Khokha MK, Hsu D, Brunet LJ, Dionne MS, Harland RM. Gremlin is the BMP antagonist required for maintenance of Shh and Fgf signals during limb patterning. Nat Genet. 2003;34(3):303–307.

51. Fenwick MA, Mansour YT, Franks S, Hardy K. Identification and regulation of bone morphogenetic protein antagonists associated with preantral follicle development in the ovary. Endocrinology. 2011:in press.

52. Cimino I, Casoni F, Liu X, Messina A, Parkash J, Jamin SP, Catteau-Jonard S, Collier F, Baroncini M, Dewailly D, Pigny P, Prescott M, Campbell R, Herbison AE, Prevot V, Giacobini P. Novel role for anti-Mullerian hormone in the regulation of GnRH neuron excitability and hormone secretion. Nat Commun. 2016;7:10055.

53. Gilbert SB, Roof AK, Rajendra Kumar T. Mouse models for the analysis of gonadotropin secretion and action. Best Pract Res Clin Endocrinol Metab. 2018;32(3):219–239.

54. Couse JF, Korach KS. Estrogen receptor null mice: what have we learned and where will they lead us? Endocr Rev. 1999;20(3):358–417.

55. Welt CK, Pagan YL, Smith PC, Rado KB, Hall JE. Control of follicle-stimulating hormone by estradiol and the inhibins: critical role of estradiol at the hypothalamus during the luteal-follicular transition. J Clin Endocrinol Metab. 2003;88(4):1766–1771.

56. Woodruff T, Krummen L, McCray G, Mather J. *In situ* ligand binding of recombinant human [^125^I] activin-A and recombinant human [^125^I] inhibin-A to the adult rat ovary. Endocrinology. 1993;133(6):2998–3006.

57. Woodruff TK, Besecke LM, Groome N, Draper LB, Schwartz NB, Weiss J. Inhibin A and inhibin B are inversely correlated to follicle-stimulating hormone, yet are discordant during the follicular phase of the rat estrous cycle, and inhibin A is expressed in a sexually dimorphic manner. Endocrinology. 1996;137(12):5463–5467.

58. Visser JA, de Jong FH, Laven JS, Themmen AP. Anti-Mullerian hormone: a new marker for ovarian function. Reproduction. 2006;131(1):1–9.

59. Weenen C, Laven JS, Von Bergh AR, Cranfield M, Groome NP, Visser JA, Kramer P, Fauser BC, Themmen AP. Anti-Mullerian hormone expression pattern in the human ovary: potential implications for initial and cyclic follicle recruitment. Mol Hum Reprod. 2004;10(2):77–83.

60. Convissar S, Armouti M, Fierro MA, Winston NJ, Scoccia H, Zamah AM, Stocco C. Regulation of AMH by oocyte-specific growth factors in human primary cumulus cells. Reproduction. 2017;154(6):745–753.

61. Dewailly D, Andersen CY, Balen A, Broekmans F, Dilaver N, Fanchin R, Griesinger G, Kelsey TW, La Marca A, Lambalk C, Mason H, Nelson SM, Visser JA, Wallace WH, Anderson RA. The physiology and clinical utility of anti-Mullerian hormone in women. Hum Reprod Update. 2014;20(3):370–385.

62. Briley SM, Jasti S, McCracken JM, Hornick JE, Fegley B, Pritchard MT, Duncan FE. Reproductive age-associated fibrosis in the stroma of the mammalian ovary. Reproduction. 2016;152(3):245–260.

63. Grgurevic L, Christensen GL, Schulz TJ, Vukicevic S. Bone morphogenetic proteins in inflammation, glucose homeostasis and adipose tissue energy metabolism. Cytokine Growth Factor Rev. 2016;27:105–118.

64. Vukicevic S, Grgurevic L. Bone morphogenetic proteins in inflammation. In: Parnham M, ed. Encyclopedia of Inflammatory Disease. Basel: Springer Verlag; 2015:1–15.

65. Lakhal B, Laissue P, Braham R, Elghezal H, Saad A, Fellous M, Veitia RA. A novel BMP15 variant, potentially affecting the signal peptide, in a familial case of premature ovarian failure. Clin Endocrinol (Oxf). 2009;71(5):752–753.

66. Kumar R, Alwani M, Kosta S, Kaur R, Agarwal S. BMP15 and GDF9 Gene Mutations in Premature Ovarian Failure. J Reprod Infertil. 2017;18(1):185–189.

67. Fonseca DJ, Ortega-Recalde O, Esteban-Perez C, Moreno-Ortiz H, Patino LC, Bermudez OM, Ortiz AM, Restrepo CM, Lucena E, Laissue P. BMP15 c.-9C>G promoter sequence variant may contribute to the cause of non-syndromic premature ovarian failure. Reprod Biomed Online. 2014;29(5):627–633.

68. Beau I, Renault L, Clemence D, Patino L, Delemer B, Laissue P, Young J, Binart N. Bone morphogenetic protein receptor variants: A new cause of primary ovarian insufficiency. Journal of the Endocrine Society. 2019;3.

69. Faure MO, Nicol L, Fabre S, Fontaine J, Mohoric N, McNeilly A, Taragnat C. BMP-4 inhibits follicle-stimulating hormone secretion in ewe pituitary. J Endocrinol. 2005;186(1):109–121.

70. Bazina M, Vukojevic K, Roje D, Saraga-Babic M. Influence of growth and transcriptional factors, and signaling molecules on early human pituitary development. J Mol Histol. 2009;40(4):277–286.

71. Sallon C, Faure MO, Fontaine J, Taragnat C. Dynamic regulation of pituitary mRNAs for bone morphogenetic protein (BMP) 4, BMP receptors, and activin/inhibin subunits in the ewe during the estrous cycle and in cultured pituitary cells. J Endocrinol. 2010;207(1):55–65.

72. Cameron V, Nishimura E, Mathews L, Lewis K, Sawchenko P, Vale W. Hybridization histochemical localization of activin receptor subtypes in rat brain, pituitary, ovary, and testis. Endocrinology. 1994;134(2):799–808.

73. Renault L, Patino LC, Magnin F, Delemer B, Young J, Laissue P, Binart N, Beau I. BMPR1A and BMPR1B Missense Mutations Cause Primary Ovarian Insufficiency. J Clin Endocrinol Metab. 2020;105(4).

74. Gurbuz F, Desai S, Diao F, Turkkahraman D, Wranitz F, Wood-Trageser M, Shin YH, Kotan LD, Jiang H, Witchel S, Gurtunca N, Yatsenko S, Mysliwec D, Topaloglu K, Rajkovic A. Novel inactivating mutations of the DCAF17 gene in American and Turkish families cause male infertility and female subfertility in the mouse model. Clin Genet. 2018;93(4):853–859.

